# Rotation-direction-dependent regulation of ATPase inhibitory factor 1 for mitochondrial ATP synthase from atomistic simulation

**DOI:** 10.1101/2024.11.19.621789

**Authors:** Ryohei Kobayashi, Kei-ichi Okazaki

## Abstract

ATPase inhibitory factor 1 (IF_1_) is an endogenous regulatory protein for mitochondrial F_o_F_1_-ATP synthase. It blocks the catalysis and rotation of the F_1_ part by deeply inserting itself into the rotor-stator interface. Recent single-molecule manipulation experiments have elucidated that forcible rotations only in the ATP-synthesis direction eject IF_1_, rescuing F_1_ from the IF_1_-inhibited state. However, the molecular mechanism of the rotation-direction-dependent process at an atomic resolution is still elusive. Here, we have performed all-atom molecular dynamics (MD) simulations of the IF_1_-bound F_1_ structure with a torque applied to the rotor γ subunit. In the torque-applying simulations, we first found that the core part of the γ subunit rotated more in response to an external torque in the synthesis direction than in the hydrolysis direction. Further rotations of the γ subunit up to 120° revealed that the conformational change of the IF_1_-bound αβ was only allowed in the synthesis direction. Also, the 120° rotation in the synthesis direction disrupted its contacts with IF_1_, destabilizing the short helix of IF_1_. After additional rotation up to the synthetic 240° state, the closed-to-open conformational change of the IF_1_-bound β subunit pulled IF_1_ outwardly, deforming the long helix of IF_1_. These stepwise destabilizations of the IF_1_ helices should be crucial for IF_1_ ejection. Our simulations also provided insight into the nullification mechanism of the hydrolytic rotation, highlighting the steric clash between F22 of IF_1_ and the β_TP_ subunit. Finally, we discussed a sufficient proton motive force to rescue F_o_F_1_-ATP synthase from the IF_1_-inhibited state.

**Significance statement:** F_o_F_1_-ATP synthase is crucial for cellular energy production. However, its reverse reaction as an ATPase can be harmful, particularly in mammalian cells. This study reveals how ATPase inhibitory factor 1 (IF_1_) controls the catalysis of mitochondrial ATP synthase and provides the atomic-level description of the rotation-direction-dependent regulation using molecular dynamics simulation. ATP-synthetic γ rotation dissociates IF_1_ from the complex, thereby recovering active ATP synthesis. However, IF_1_ completely suppresses the γ rotation through multiple mechanisms to prevent wasteful ATP consumption in ATP-hydrolysis mode. Our analysis also reveals that sufficient torque in the synthesis direction is required to rescue F_o_F_1_-ATP synthase from the IF_1_-inhibited state. These findings offer valuable insights for developing inhibitors by leveraging the structural and functional similarities with IF_1_.

## Introduction

F_o_F_1_-ATP synthase (F_o_F_1_) is the terminal enzyme of oxidative phosphorylation, which synthesizes ATP from ADP and inorganic phosphate (P_i_) driven by the electrochemical proton gradient, or the proton motive force (*pmf*) (1–3). F_o_F_1_ represents two rotary motors, the membrane-embedded F_o_ domain and the water-soluble F_1_ domain, which are connected by a rotary shaft that efficiently transmits the torque. F_o_F_1_ is a reversible motor, where the rotary motion of the central stalk is switched upon the balance of the free energy of ATP hydrolysis versus *pmf* across the biological membranes (4). F_o_F_1_ works as an ATP synthesis motor when enough *pmf* is maintained: proton translocation through F_o_ induces the rotation in the clockwise (CW) direction, viewed from the outside of the membrane. In the hydrolysis mode, where *pmf* is low, F_1_ hydrolyzes ATP to rotate the rotor complex counter-clockwise (CCW) to pump protons. Since the physiological role of F_o_F_1_ is ATP synthesis, several regulator proteins prevent the wasteful ATP hydrolysis by F_1_, which is detrimental to the cell. Bacterial ATP synthase has an endogenous inhibitor called ε subunit intrinsically bound to the γ subunit (5–9). Mammalian ATP synthase has ATPase inhibitory factor 1 (IF_1_) as a principal regulator for ATP hydrolysis (10–14). The inhibition must be released when *pmf* is recovered so that F_o_F_1_ can synthesize ATP again.

F_1_ comprises the stator ring and the rotary shaft (15). The rotor γ subunit is inserted into the central cavity of the stator α_3_β_3_-ring. The catalytic sites for ATP synthesis and hydrolysis are located at the αβ interfaces, mainly on the β subunit, although some residues in the α subunit are also involved. The resolved structures have revealed the conformational change of three β subunits with different nucleotide states, whereas the α subunit took almost the same conformation (16–18). Typically, two of the three β subunits have the bound nucleotides; one for ATP analog (called β_TP_), the other for ADP (β_DP_). They adopt a closed conformation in which the C-terminal domain of the β subunit swings inwardly. The other β subunit (β_E_) has no bound nucleotide and takes an open conformation with its C-terminus staying away from the γ subunit. The open-to-closed conformational change upon the nucleotide binding is essential for the chemo-mechanical coupling of F_1_. Furthermore, the crystal structures of IF_1_ bound F_1_ provide insight into the inhibition mechanism (19, 20). IF_1_ forms two distinctive α-helices, the short helix (residues 14-18) and the long helix (residues 21-50). The N-terminus of IF_1,_ including the short helix, is inserted near the γ subunit, whereas the long helix of IF_1_ is bound to the α_DP_β_DP_ interface, mainly the C-terminus of the β_DP_ subunit.

Our recent single-molecule manipulation experiments clarified the rotation-direction-dependent activation from the IF_1_ inhibition (21). The IF_1_-inhibited F_1_ was efficiently activated by the forcible CW rotation (the synthesis direction) of the γ subunit, although the CCW rotation (the hydrolysis direction) did not have any effect. The stall-and-release experiments provided more information about the angle-dependent manner of F_1_ activation from the IF_1_ inhibition. The forcible CW rotation with more than 200° enhanced the activation probability, suggesting that IF_1_ ejection is coupled to the ATP synthesis reaction. Furthermore, the N-terminus-truncated mutants of IF_1_ elucidated the importance of F22 in IF_1_: ΔIF_1_(1-22) lost the asymmetric activation feature, whereas ΔIF_1_(1-19) still maintained the feature. These experiments significantly contribute to an understanding of the IF_1_ inhibition. However, due to the spatial resolution of the experiments, detailed molecular mechanisms have remained elusive. Molecular dynamics (MD) simulations with an external force mimicking the single-molecule manipulation forces can complement the experimental observations and provide atomic-level descriptions of the underlying events (22, 23). MD simulations have also clarified the mechano-chemical coupling of F_1_ in the absence of IF_1_ with a torque applied on the γ subunit (24–29).

In this paper, we first performed structural analysis to characterize the IF_1_ bound structures. Then, we have performed atomistic MD simulations of the bovine mitochondrial F_1_ (*b*MF_1_)-IF_1_ complex to clarify the molecular mechanism of the rotation-direction-dependent inhibition of IF_1_. To examine the impact of the rotation of the γ subunit to IF_1_-bound F_1_, the γ subunit was forcibly rotated in the CCW and CW directions by the previously developed flexible rotor method. We analyzed the relative conformational change of the IF_1_-bound αβ pair and short helix deformation at the N-terminus of IF_1_.

Then, we further rotated the γ subunit to the angle at which the open-to-closed conformational change of the IF_1_-bound β subunit would be assumed and analyzed the long helix deformation of IF_1_. We also examined how IF_1_ inhibits the CCW rotation, focusing on the β_TP_ subunit adjacent to the IF_1_-bound αβ pair.

## Results

### Structural characterization of IF_1_-bound *b*MF_1_ structures

The crystal structure of *b*MF_1_ with a monomeric form of IF_1_ (IF_1_^1-60^) provides initial insight into the inhibitory complex (PDB ID: 2v7q) (19). IF_1_ possesses two distinctive α-helices, the short helix (residues 14-18) and the long helix (residues 21-50). The N-terminus of IF_1,_ including the short helix, is inserted near the γ subunit, whereas the long helix of IF_1_ is bound to the α_DP_β_DP_ interface, mainly the C-terminus of the β_DP_ subunit. Following this structure, the crystal structure with three IF_1_s bound to each αβ interface, referred to as *b*MF_1_-(IF_1_)_3_, was reported (PDB ID: 4tt3) (20). The *b*MF_1_-(IF_1_)_3_ structure showed a stepwise folding of IF_1_ upon binding to the catalytic αβ subunit of F_1_. In IF_1_ bound to αβ_E_, only the second half of the long helix (residues 32-49) was resolved. In IF_1_ bound to αβ_TP_, the whole long helix (residues 23-50) was resolved. IF_1_ bound to αβ_DP_ adopted the most folded state, i.e., residues 11-50 were resolved. The partial folding forms of IF_1_ at αβ_E_ or αβ_TP_, observed in other structures with two or three IF_1_s (PDB IDs: 4tsf, 4tt3, 4z1m) (20, 30), were thought to represent intermediates to the fully-inhibited state. IF_1_ at αβ_DP_ always showed the fully-folded state (PDB IDs: 1ohh, 2v7q, 4tsf, 4tt3, 4z1m) (19, 20, 30, 31), where the short helix in the N-terminus was resolved as well as the long helix in the C-terminus.

We characterized the IF_1_-bound *b*MF_1_ structures among all available F_1_ structures. To identify relative conformations of the IF_1_-bound αβ, we have performed principal component analysis (PCA) on the protein structures, transforming the high-dimensional data into a few dimensions. Although the fundamental aspects of the F_1_ structures have already been reported (32), we included the crystal structures published since the last report. The currently available 26 *b*MF_1_ structures from the Protein Data Bank (PDB) were analyzed. Each F_1_ structure contains three αβ pairs with different conformations and nucleotide states, resulting in a total of 78 αβ pairs. Based on the orientation of the γ subunit, three αβ pairs were named αβ_E_, αβ_TP_, and αβ_DP_, respectively. Among them, IF_1_ binds to 10 αβ pairs (1ohh, 2v7q, 4tsf, 4tt3, 4z1m in αβ_DP_, 4tt3, 4z1m in αβ_TP_, and 4tsf, 4tt3, 4z1m in αβ_E_).

Here, we analyzed Cα atoms of the αβ pairs resolved in all structures (see *Methods*). The resulting top two PCA modes explained 97% of the total motions, with the first principal component (PC1) representing the opening/closing motion of the β subunit and the second principal component (PC2) representing the loosening/tightening motion at the αβ interface. This analysis has well captured the representative conformational motions of the αβ pairs during catalysis. The PC1-PC2 plot in Fig. 1B shows some dense clusters, three of which represent the typical conformational states of αβ_E_, αβ_TP_, and αβ_DP_ (circled by the dashed lines). αβ_E_ adopts an open conformation of the β subunit, αβ_TP_ and αβ_DP_ adopt a closed conformation with its interface loose or tight, respectively. The IF_1_-bound αβ_E_ in 4tsf, 4tt3, 4z1m, and the αβ_TP_ in 4tt3, 4z1m were also included in these αβ_E_ and αβ_TP_ clusters, respectively, suggesting that the partially-folded IF_1_ does not affect the conformational states of αβ. The other cluster (circled by the solid line) contained the five IF_1_-bound αβ_DP_’s, showing an intermediate PC2 value between the αβ_DP_ and αβ_TP_ with a similar PC1 value. This result indicates that the αβ interface is not fully tightened as the typical αβ_DP_ cluster, while the β subunit takes an almost closed form (Fig. 1B and 1C). This incomplete closure of the αβ interface is one of the distinct features that discriminate the fully-folded IF_1_-bound state from the typical αβ_DP_ state, as seen in the PC2 value.

**Fig. 1.**
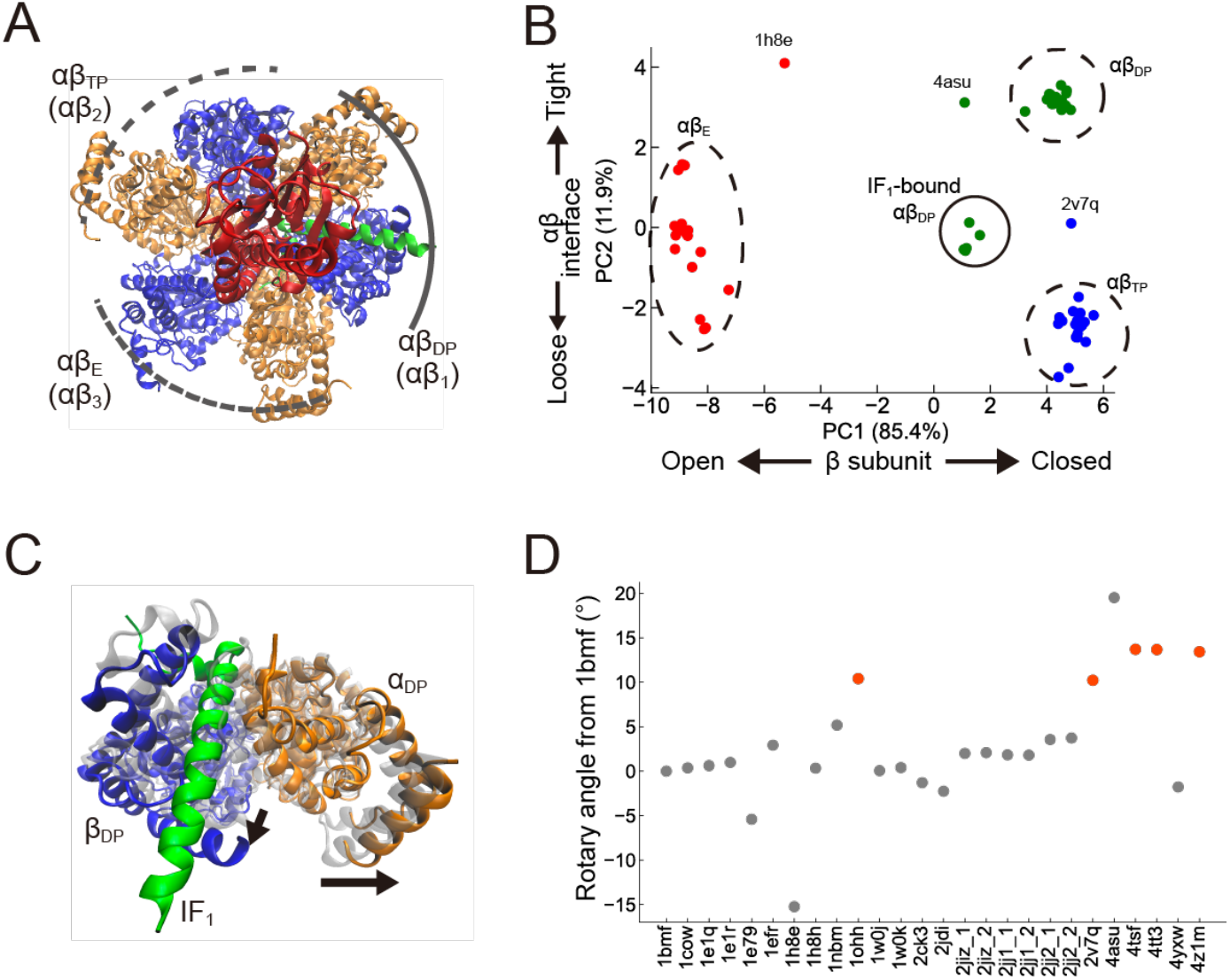
Characterization of IF_1_-inhibited structures. (A) The top view of the F_1_-IF_1_ structure (PDB: 2v7q). The α, β, γ, and IF_1_ are shown in orange, blue, red, and green, respectively. The δ and ε subunits are omitted in this figure. (B) Principal Component Analysis (PCA) of αβ pairs. The 78 αβ pairs from 26 *b*MF_1_ structures were projected on PC1 and PC2. The nucleotide state of each αβ pair, Empty, TP, and DP, are shown in red, blue, and green, respectively. The values in the parentheses of the axis labels refer to the 1^st^ and 2^nd^ eigenvalue contributions. The units of each axis are in nanometers (nm). (C) Structural comparison of the αβ_DP_ in the IF_1_-inhibited structure (PDB: 2v7q, orange, blue, and green) with the ground-state structure (PDB: 2jdi, gray/transparent). Arrows represent the interface motion of IF_1_-bound αβ_DP_ compared to the catalysis-waiting state. (D) The rotary angle of the γ subunit in 26 *b*MF_1_ structures, where that of the 1bmf structure was defined as 0 degree. The orange points represent the IF_1_-bound structures.

We also quantified the rotary angle of the γ subunit in all 26 structures from that of the 1bmf structure. The rotary angle was determined by aligning the α_3_β_3_ of all F_1_ complexes and then rotating each γ subunit to best fit the 1bmf structure (see *Methods*). Fig. 1D shows a clear difference between the IF_1_-bound structures and the others: the γ subunit of the IF_1_-bound structures was rotated +10°-15° in the hydrolysis (CCW) direction. This analysis is consistent with the stall position of IF_1_ inhibition observed in the single-molecule experiment^21^: IF_1_-stall positions were 90° from the ATP-binding waiting position, whereas the catalysis-waiting positions were 80°. The PCA and γ angle results indicate that the IF_1_-bound structures are qualitatively different from the other *b*MF_1_ structures representing the catalysis-waiting state.

### Torque-applying simulations in CW and CCW directions

In our single-molecule manipulation experiments, forcible CW rotation (the synthesis direction) of the γ subunit by magnetic tweezers led to F_1_ activation from the IF_1_ inhibition. In contrast, activation was not observed in CCW rotation (the hydrolysis direction) (21). To explore the molecular basis of this rotation-direction-dependent activation, we have performed an all-atom, explicit-solvent MD simulation from the IF_1_-bound F_1_ structure (PDB ID: 2v7q). The γ subunit was forcibly rotated in the CCW and CW directions by the previously developed flexible rotor method (27, 28), implementing the realistic and adaptive rotations of the γ subunit with only the average angle controlled (Fig. 2A and 2B). To see the rotation-direction dependence in or near the IF_1_-inhibited position, we first conducted 40° torque simulations with the angular velocity of *ω* = 1°/ns, applying various force constants, κ = 10^4^, 10^5^, 10^6^ kcal·mol^-1^·rad^-2^ (Fig. 2C-2F). Note that 10^5^ kcal·mol^-1^·rad^-2^ ∼ 30.5 kcal· mol^-1^ · deg^-2^ was used in the previous study (28). The γ subunit shows non-uniform rotations, particularly between the residues buried inside and outside the α_3_β_3_-ring (Fig. 2A and 2B). With weaker restraints, the core part of the N-terminus (residue 1-26) and the C-terminus (residues 228-272) of the γ subunit was significantly lagged compared to the protruded part. It represents the structural flexibility of the γ subunit, which is supported by the root mean squared deviation (RMSD) analysis (Fig. S1), where the γ subunit was twisted with a weaker restraint. In both directions, a larger value κ resulted in a more significant rotation of the core part of the γ subunit (Fig. 2C). Overall, the essential characteristics of the current simulation system of the IF_1_-bound F_1_ were similar to that reported in the earlier studies (28).

**Fig. 2.**
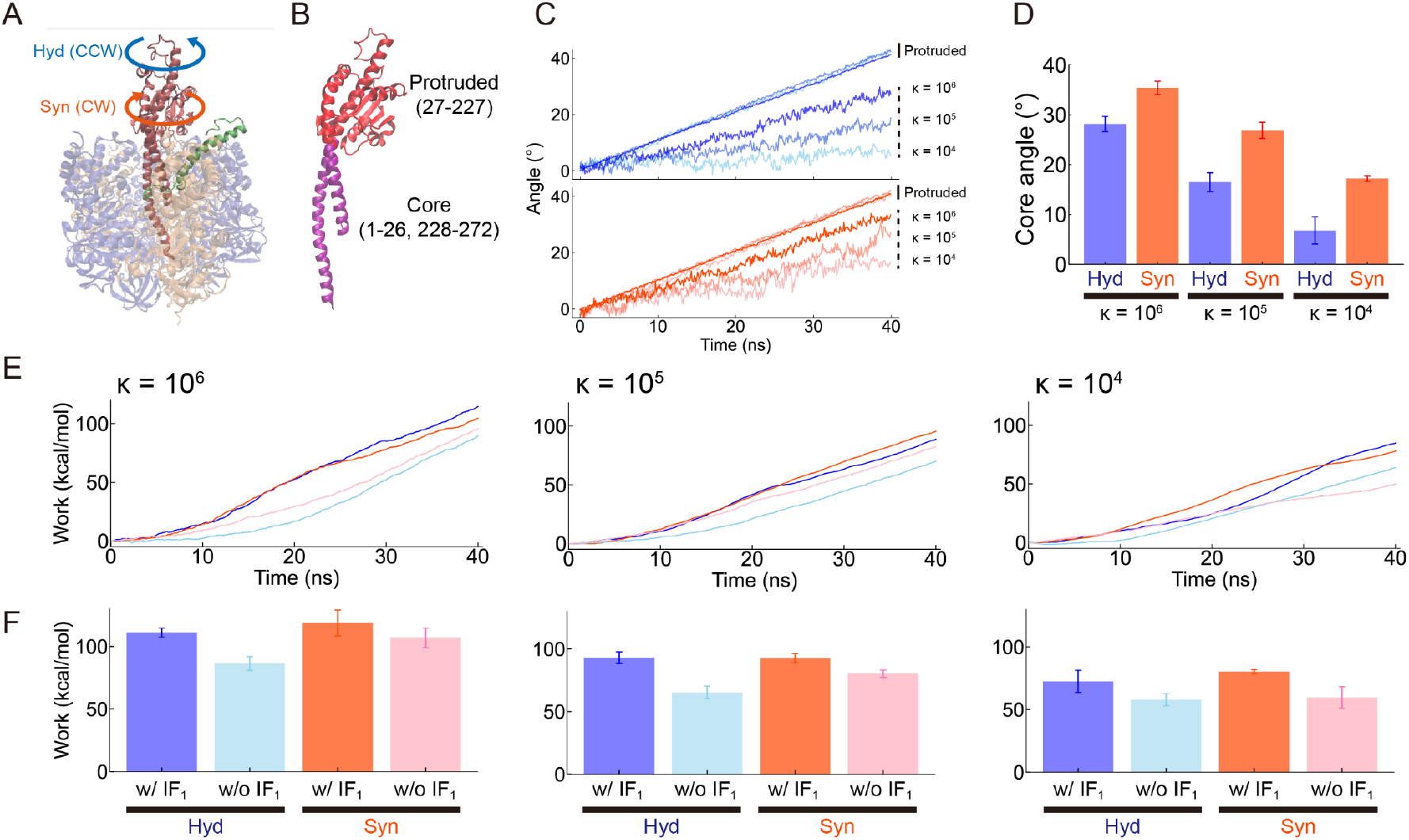
The 40° torque simulation with various force constants. (A) Overview of the torque-applying simulation. The γ subunit was rotated in hydrolysis (counter-clockwise; CCW) or synthesis (clockwise; CW) direction at ω = 1 °/ns. (B) The structure of the γ subunit. The core part (residues 1-26 and 228-272) is colored purple, and the protruded part (residues 27-227) is colored red. (C) The γ subunit rotation of the core part and the protruded part. The protruded and core angles show linear and jiggling behaviors, respectively. The core angles are shown in gradation concerning different force constants. (D) The final core angles from the 40 ns simulations. The mean values and SD error bars are calculated from 3 independent simulations. (E) Work profiles over the simulation time. Blue and orange lines represent the simulation with IF_1_, and cyan and pink lines represent IF_1_-free simulations. (F) The final work from the simulations. The mean values and SD error bars are calculated from 3 independent simulations.

One of the most notable findings in our simulations is that at a given value of κ, the core of the γ subunit rotated more in the synthesis direction than in the hydrolysis direction (Fig. 2D). The preference was more evident with κ = 10^4^ kcal·mol^-1^·rad^-2^, although the same trend was also seen with larger κ. A large κ may allow the system to overcome slight differences upon rotation, such as molecular friction and structural asymmetry. However, these subtle differences become more pronounced with a weaker κ, resulting in a factor of two differences in the cumulative rotation angle. We also tested the IF_1_-free *b*MF_1_ simulation and found a slightly biased rotation towards the hydrolysis direction on the contrary (Fig. S2). This is consistent with the previous single-molecule experiment (33), the theoretical study (34), and the computational analysis (28): the phosphate release in the β_E_ promotes CCW (hydrolysis) rotation of γ subunit from the catalytic dwell state. These findings suggested that the asymmetrical mobility of the γ subunit towards the CW (synthesis) direction distinguishes the IF_1_-inhibited state from the active catalytic state. To further quantify the differences between these two states, we have measured the non-equilibrium work during the forcible rotations of the γ subunit (Fig. 2E and 2F, and Fig. S3; *Methods*). The total work required for 40° rotations exhibited a clear trend at any κ: the IF_1_-inhibited state required 20% more work than the typical catalytic dwell state (Fig. 2F). We also estimated the rotary torque during the simulation (Fig. S4). Since the torque plot over the simulation time showed large fluctuations, we calculated the average torque values over the simulations. The results were consistent across all the simulations: simulations with IF_1_ required more torque than the IF_1_-free simulations. The estimated torque values were 2-4 times larger than the experimental values reported in the single-molecule experiments, 40 pN·nm/rad. Overall, these results reflect the mechanical stiffness of the IF_1_-inhibited state compared to the catalysis-waiting state. Further discussion will be provided in the *Discussion* section.

### 120° rotation in CW and CCW

To further explore the rotation-direction-dependent activation observed in the single-molecule manipulation experiment, we rotated the γ subunit up to 120° in both directions. A force constant, κ = 10^6^ kcal·mol^-1^·rad^-2^, was applied in the following simulations, mimicking the single-molecule manipulation experiment with extremely greater force than that F_1_ generates. This simulation setup enabled a 100° rotation in the core part with an average 120° rotation of the whole. It should also be noted that, in the later sections of this paper, the γ subunit was rotated more than 120° from the initial state, which changes the relative position of each αβ to the orientation of the γ subunit. To avoid misunderstandings regarding the name of each αβ, we rename αβ_DP_ (bound to IF_1_), αβ_TP_, and αβ_E_ in the initial structure to αβ_1_, αβ_2_, and αβ_3_, respectively (Fig. 1A).

To analyze the conformational changes of three αβs, we projected the simulation trajectories onto the PC1 and the PC2 (Fig. 3A for the αβ_1_ and Fig. S5A for the αβ_2_ and the αβ_3_, respectively). During CW rotation (the synthesis direction) of the γ subunit, the PC2 of the αβ_1_ decreased down to the level of the αβ_TP_ cluster (Fig. 3A, left), suggesting that the IF_1_-bound αβ interface was loosened upon the rotation. We note that such a loosening motion of the IF_1_-bound αβ was not observed in torque-free simulations (Fig. S5B). In contrast, CCW rotation (the hydrolysis direction) did not induce any appropriate conformational change of the IF_1_-bound αβ (Fig. 3A, right). This clear dependence upon the rotary direction was also observed in Fig. 3B, where the relative amount of accomplished conformational change from the initial IF_1_-inhibited state to the next catalytic state is plotted. These results indicated that the rotation-direction-dependent activation observed in the single-molecule manipulation experiments originates from whether the IF_1_-bound αβ can appropriately change its conformation upon rotation of the γ subunit.

**Fig. 3.**
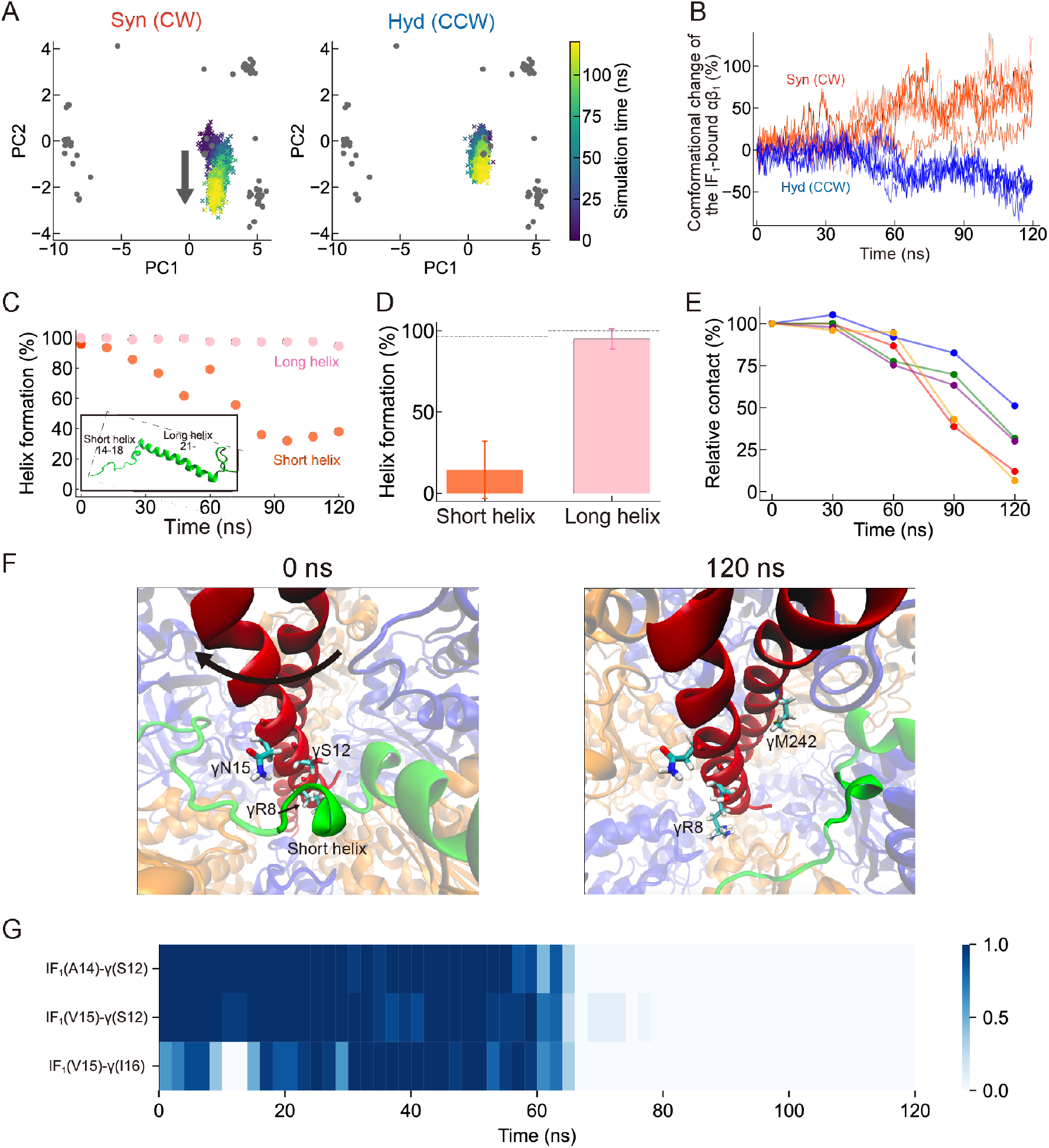
The 120° torque simulation. (A) Conformational change of the IF_1_-bound αβ pair upon the CW (left) and CCW (right) rotation, projected onto the PC1-PC2 plane. The gray dots correspond to the X-ray crystal structures (Fig. 1B). (B) The relative amount of accomplished conformational change of the IF_1_-bound αβ_1_ pair upon the γ rotation quantified from the PC2 value. Different shades of color represent five independent trajectories. (C) A typical example of the short helix (residues 14-18) and the long helix (residues 21-45) formation during the CW 120° simulation. Helix formation at time 0 was calculated from the simulation result before the torque-applying simulation. Inset is the initial structure of IF_1_. (D) Average helix formation in the final 10 ns (110-120 ns) of the CW simulation. The dotted lines represent the value calculated from the simulation result before the torque-applying simulation. (E) Time-dependent changes in contacts between the short helix and the γ subunit during the CW simulation. The relative contact values were calculated by setting the contact number from the equilibrium simulation as 100%. The data points at 30 ns, 60 ns, 90 ns, and 120 ns were calculated from the number of contacts in the preceding 30 ns interval. (F) The snapshots at 0 ns (*left*) and 120 ns (*right*) during the CW rotation. The bright green region represents the short helix of IF_1_, while the rest is shown as transparent. Representative residues are shown in both figures, γR8, γS12, and γN15. γM242 from the C-terminus is shown in the right figure, which approaches the short helix at the end of some simulations. (G) The time-dependent contact changes of the specific residue pairs between the short helix and the N-terminus of γ subunit; IF_1_(A14)-γ(S12), IF_1_(V15)-γ(S12), IF_1_(V15)-γ(I16). Dark blue represents a high contact ratio, while light blue represents a low one. The same simulation trajectory was used to illustrate Fig. 3C, 3F, and 3G.

To investigate the impact of the 120° rotation on the IF_1_ conformation, we have quantified the helix formation of IF_1_ during the CW (synthesis) rotation. The most folded form of IF_1_ possesses two helices linked by a glycine kink: the short helix of residues 14-18 that interacts with the γ subunit and the long helix of residues after 21 that principally interacts with the β_DP_ subunit (Fig. 3C, inset). Five independent simulations showed that the short helix was gradually deformed during the γ rotation, whereas the long helix was almost intact (Fig. 3C and 3D). We also analyzed the contact of the amino acid residues between the short helix and the γ subunit (Fig. 3E and 3F). In the torque-free simulation, the N-terminal residues of the γ subunit, such as R8, R9, K11, S12, N15, and I16, were in contact with the short helix. However, in the last part of the torque-applying simulation, where the γ subunit was sufficiently rotated to the CW direction, these residues moved far from the short helix. The total number of contacts significantly reduced, although a few residues have formed new contacts, including the C-terminus of the γ subunit such as γM242 (the right panel in Fig. 3F). The time-dependent contact change between some specific amino acid pairs is also along this contention (Fig. 3G). These results suggest that loss of the contacts with the γ subunit upon rotation leads to the destabilization of the short helix.

### Conformational transition of β at CW 240° induces deformation of the long helix

The previous single-molecule stall-and-release manipulation experiment showed a remarkable increase in the activation probability at CW 200°-240°, suggesting that IF_1_ is ejected through the closed-to-open conformational transition of the β subunit upon ATP release synthesized from ADP and P_i_ (21). Thus, implementing conformational transitions in our simulations should be crucial for observing IF_1_ ejection. Indeed, our CW-120°-rotation simulation showed a partial, though not complete, conformational change of all three αβs. It is also challenging to go beyond 120° rotation because the nucleotide state of each αβ has to change. To reflect the nucleotide state change in our simulations, we switched the nucleotide states of the structure after 120° rotation, replacing the initial nucleotide states with the ones for the CW 120° state (see *Methods*). After relaxation, we performed a targeted MD simulation to complete the conformational change for each αβ at 120°. As a result, αβ_1_ (bound to IF_1_), αβ_2_, and αβ_3_ reached the TP, Empty, and DP state, respectively. Then, the second round of the torque-applying simulation was performed to forcibly rotate γ subunit from the CW 120° to the CW 240° state. During the targeted MD and the subsequent forcible γ rotation, no significant conformational changes of IF_1_ were observed, although αβs showed conformational changes (Fig. S6A).

After CW 240° rotation, the nucleotide states were again adjusted to the corresponding bound nucleotide states at this angle (Fig. 4A). The targeted MD completed conformational changes across all αβs, particularly the closed-to-open conformational change of the IF_1_-bound β_1_ subunit, as seen in the PC1 value (Fig. S6B). During 50 ns of the targeted MD, the long helix showed significant conformational change from a linear to a bent structure around its middle part (Fig. 4B and 4D). The secondary structure analysis revealed the deformation of the long helix (Fig. 4C), though the extent of deformation varied among the four independent simulations (Fig. S7). In two simulations, the long helix underwent severe disruption near the short helix (residues 21-30), similar to the partially-resolved form of IF_1_ in crystal structures (see *Discussion* for more details). The other two simulations exhibited a loss of stability around the middle part of the long helix (residues 31-37). Overall, our simulations underscored a considerable destabilization of the long helix, which had been remarkably stable in earlier stages of the simulations described in this study. These structural dynamics were mechanistically linked to the swinging motion of the C-terminus of the β subunit, which pulls the C-terminal of IF_1_ (residues L42-L45) outwardly. Notably, the bending motion of IF_1_ was not observed in equilibrium simulations, in which no conformational change of αβ pairs was observed. These deformations of IF_1_ also weakened the salt bridge between E30 of IF_1_ and R408 of IF_1_-bound β (β_1_), which is a well-known interaction to stabilize the inhibitory complex (Fig. 4E). Given that the E30A mutant deficient in this salt bridge resulted in an unstable state in solution experiments (35, 36), these observations likely represent an intermediate state in the process of the dissociation of IF_1_ from F_1_. Altogether, our simulations revealed that the closed-to-open conformational change of the IF_1_-bound β_1_ subunit significantly destabilized the long helix, which would lead to the IF_1_ dissociation.

**Fig. 4.**
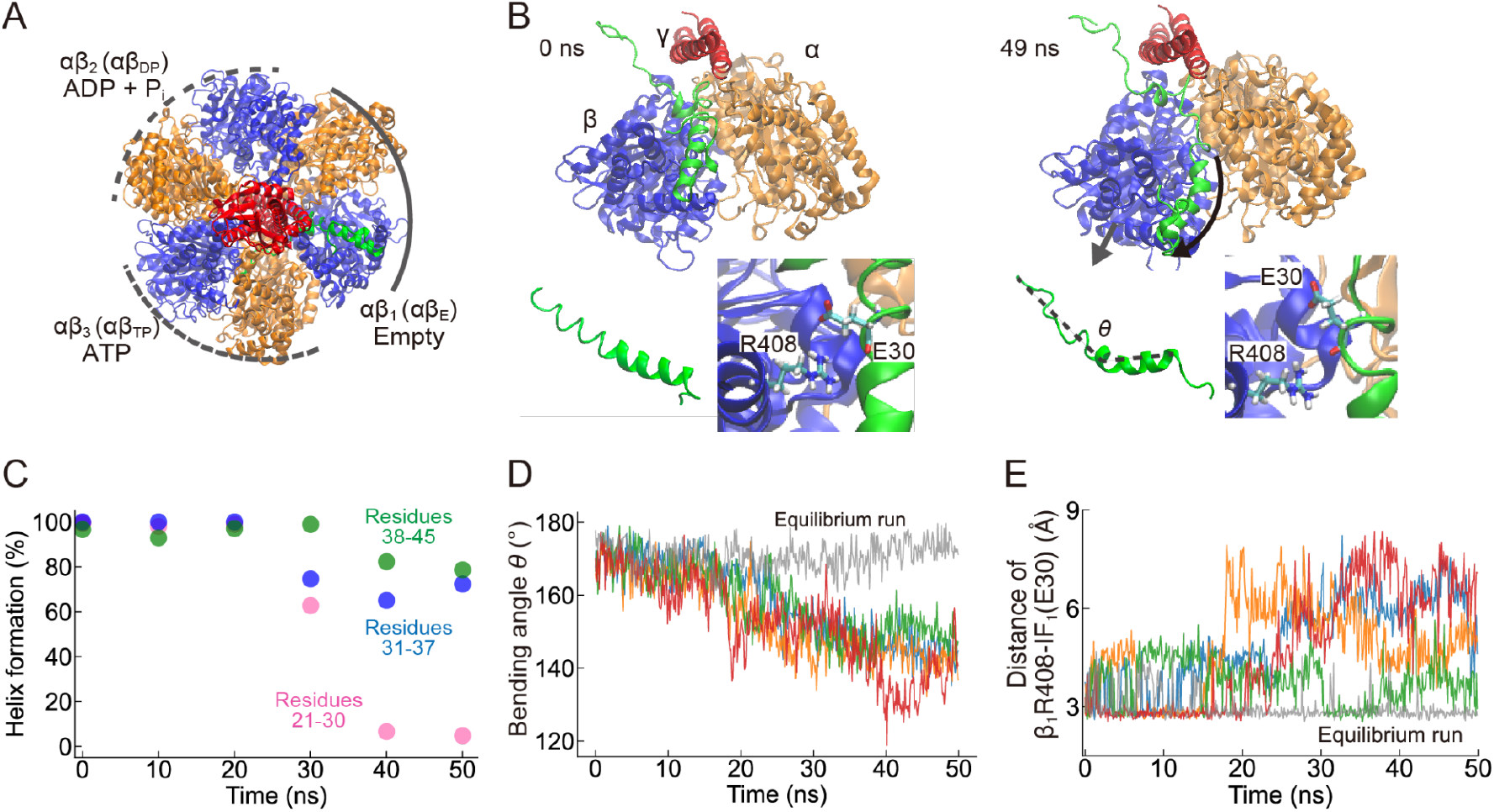
Deformation of the long helix at CW 240°. (A) Overview of the CW 240° state, where the bound nucleotides are also shown. (B) Snapshots from the targeted MD at 0 ns (*Left*) and 49 ns (*Right*). The inset in the left lower in each panel is IF_1_, where residues 21-50 are shown. The inset in the right lower in each panel is the enlarged view of the R408 of IF_1_-bound β (β_1_) and E30 of IF_1_. (C) A typical example of the long helix (residues 21-45) deformation. The long helix was divided into three parts: residues 21-30 (pink), 31-37 (blue), and 38-45 (green). Helix formation at time 0 was estimated from the simulation result before the targeted MD. (D) The bending angle of the long helix during the targeted MD, which was defined by the angle formed by the Cα atoms of residues 21, 38, and 45. The gray line represents the result of the equilibrium run before the targeted MD. (E) The minimum distance between E30 of IF_1_ and R408 of IF_1_-bound β (β_1_). The gray line represents the result of the equilibrium run before the targeted MD.

### Nullification of CCW rotation

So far, we have described that the forcible CW rotation (the synthesis direction) of the γ subunit destabilizes and eventually ejects IF_1_ through the short and the long helices deformation. Another fundamental question regarding IF_1_ regulation is how IF_1_ halts or nullifies the CCW rotation (the hydrolysis direction) at the atomic level. The forcible CCW rotation never activated F_1_ from the IF_1_ inhibition in the experiment. That is, it is completely nullified. To further explore this question, we revisit and analyze the forcible 120° rotation simulations, paying particular attention to the CCW direction. During the CCW 120° rotation, the short helix was partially destabilized (Fig. 5A). Still, it was less affected compared to the CW rotation (Fig. 3E). This result indicated that the asymmetric change in the stability of the short helix occurred in a rotation-direction-dependent manner. Additional analysis confirmed that the number of contacts between the short helix and the γ subunit was less affected during the CCW rotation (Fig. 5B). This is because the N-terminus of γ, such as γK11 and γN15, moved closer to and in contact with the short helix of IF_1_ (Fig. 5C and Fig. S8), possibly preventing the abrupt disruption of the short helix.

**Fig. 5.**
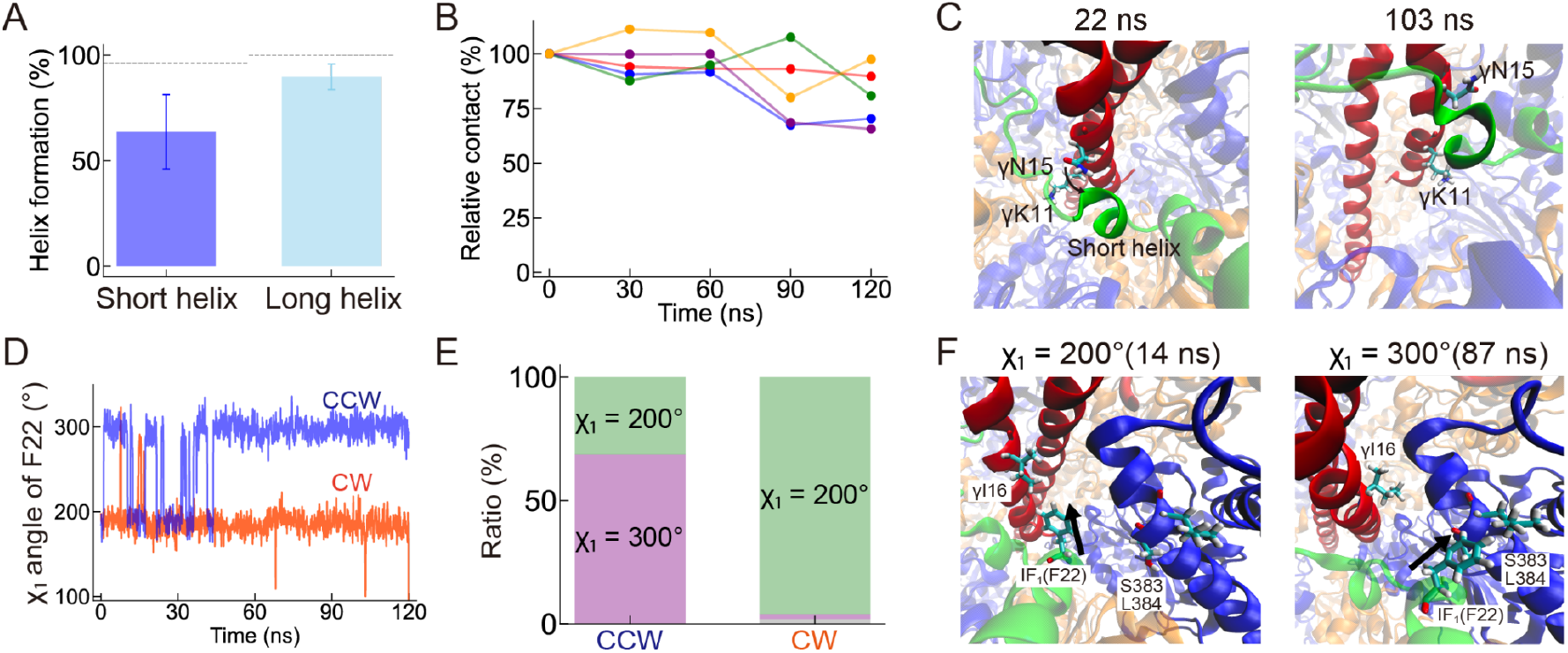
The 120° torque simulation in CCW direction. (A) Average helix formation in the final 10 ns (110-120 ns) of the CCW simulation. The dotted lines represent the equilibrium value calculated from the simulation before applying torque. (B) Time-dependent contact changes between the short helix and the γ subunit during the CCW simulation. See also Fig. 3D and 3E for reference of the CW rotation. (C) Snapshots focusing on the short helix of IF_1_ and the γ subunit. (D) The side-chain dihedral angle χ_1_ of F22 of IF_1_. The blue and orange lines represent the 120° forcible simulation in CCW and CW, respectively. (E) The ratio of χ_1_ = 200° state and the χ_1_ = 300° state in CCW and CW are shown in green and purple, respectively. The dihedral angle that was not assigned to either χ_1_ = 200° or χ_1_ = 300° is shown in gray. (F) Snapshots from the CCW simulation. The left snapshot represents the χ_1_ = 200° state, where the phenyl group of F22 in IF_1_ was oriented toward the I16 of the γ subunit. The right snapshot represents the χ_1_ = 300° state, where the phenyl group of F22 in IF_1_ was oriented toward the S383 of the β_TP_ subunit. Arrows represent the side-chain directions of F22.

The previous single-molecule manipulation experiments with N-terminal-truncated mutants of IF_1_ emphasized the importance of the short helix (21). The rotation-direction-dependent activation was lost with IF_1_(Δ1-22), although other truncated mutants, including IF_1_(Δ1-19), still maintained it. That is, F_1_ with IF_1_(Δ1-22) was activated even by the forcible CCW rotation. This finding highlights the crucial role of the residues after 19, *i*.*e*., G20, A21, and F22. Given the bulky nature of F22 among these residues, it is reasonable to assume that the interaction of F22 with F_1_ would be responsible for this feature. Along this contention, we first analyzed the contact of F22 with other subunits, but there were no apparent differences in the F22 contact between the CCW and the CW rotation (Fig. S9). Both simulations commonly increased the total contact with F_1_, particularly with the β_TP_ subunit. However, this residue’s side-chain dihedral angle χ_1_ showed a strong dependence upon the rotary directions. Two discrete states of χ_1_ = 200° and χ_1_ = 300° were identified during the CCW 120° rotation (the blue line in Fig. 5D). In contrast, the CW rotation favored the χ_1_ = 200° than the χ_1_ = 300° state (the red line in Fig. 5D). These two states resulted in different contacts with F_1_: the χ_1_ = 200° state was facing toward the γ subunit such as γI16 (Fig. 5F, left), whereas the χ_1_ = 300° state toward the neighboring β_2_ (β_TP_) subunit, including S383 and L384 (Fig. 5F, right). This means that F22 in the χ_1_ = 300° state directly pushes against β_TP_, possibly destabilizing the α_3_β_3_-ring. In contrast, χ_1_ = 200° state observed in the CW rotation did not interfere with the rotary dynamic and the structural stability of F_1_.

## Discussion

### Activation from the IF_1_-inhibited state

In this study, we have performed MD simulations of the *b*MF_1_-IF_1_ complex with forcible rotation of the γ subunit and analyzed the atomistic mechanism of the rotation-direction-dependent regulation of IF_1_. We have revealed the stepwise destabilization of IF_1_ upon the CW rotation (the synthesis direction), which is consistent with the reported crystal structures of *b*MF_1_ with two or three IF_1_ bound. In our simulations, the short helix of IF_1_ became unstable after CW 120° rotation (Fig. 3C and 3D), which well reflects the IF_1_ conformation bound to the αβ_TP_ state in the crystal structure, where the short helix was not resolved. At CW 240° after the targeted MD of the IF_1_-bound αβ changed to αβ_E_, IF_1_ showed a partially folded form of the long helix with only residues 31-49 folded (Fig. 4), as in the IF_1_ bound to the αβ_E_ state in the crystal structure. These comparisons verify our atomistic simulations upon the forcible γ rotation and the subsequent targeted MD. Furthermore, our simulations clarified the dynamic mechanisms of how the short and long helices are deformed in atomic details. While the short helix was deformed by losing contact with the γ subunit, the long helix was deformed through the closed-to-open conformational change of the IF_1_-bound β subunit.

In addition to the dynamics of the IF_1_ ejection, we also obtained an insight into the energetics of the IF_1_-inhibited state. The torque-applying simulations with various force constants found that the IF_1_-inhibited state requires 20% larger work and torque for initiating rotation than the catalysis-waiting state (Fig. 2E, 2F, Fig. S4). Although the force constant affects the work and torque values, this trend remained unchanged. The result may reflect the mechanical stiffness or molecular friction of the IF_1_-inhibited state. This finding is reasonable from the physiological viewpoints of IF_1_. IF_1_ halts the rotation of F_o_F_1_ under low *pmf* conditions, where F_1_ forcibly rotates F_o_ in the CCW direction by hydrolyzing ATP. When *pmf* returns to normal levels, F_o_F_1_ attempts to switch its rotary direction to synthesize ATP. However, a slightly greater torque for CW rotation may not be sufficient to drive efficient ATP synthesis, as even small environmental changes could cause the system to revert to CCW rotation. Therefore, sufficient torque higher than the standard value of approximately 40 pN·nm/rad would be necessary to ensure robust CW rotation. The system can eject IF_1_ and maintain stable CW rotation only once this higher torque is sustained. Assuming that *b*MF_1_ generates a torque of 40 pN·nm/rad like other F_1_s at any rotary angle, we can estimate the torque needed for ejecting IF_1_ as 48 pN·nm/rad, which is 20% larger than the standard. By equating the work done by this torque to the input *pmf* with the proton stoichiometry of 8 protons/turn in mitochondrial ATP synthase, we obtain the *pmf* of 235 mV to reactivate ATP synthesis. Considering 150-210 mV as a typical *pmf* value inside mitochondria reported in some experiments (3, 37), this estimate is slightly larger but reasonable based on the abovementioned discussion. Several studies suggested that IF_1_ inhibits catalysis not only under ATP hydrolysis but also under ATP synthesis conditions (38, 39). In these experimental conditions, a *pmf* may be sufficient for normal catalysis in IF_1_-free states but insufficient to overcome the IF_1_-inhibited state, according to the discussion above. The measurement of *pmf* required for activation from the IF_1_ inhibition can confirm this hypothesis. However, the precise control of *pmf* with reconstituted liposomes or inverted vesicles from mitochondria is experimentally challenging.

### Nullification of CCW rotation in the IF_1_-inhibited state

Our research also sheds light on how IF_1_ halts or nullifies the CCW rotation, which is not easily accessible in other studies. Compared to the ADP inhibition, which is a well-known inhibition mechanism of F_1_, the directionality of the IF_1_ inhibition is much more rigorous. The ADP inhibition only affects the hydrolysis (CCW) rotation as IF_1_ (40). However, it still allows spontaneous activation through the CCW rotation (41, 42). In contrast, the IF_1_ inhibition is never activated by the CCW rotation, literally nullifying it. To clarify the nullification mechanism, we first found that the core of the γ subunit was rotated less in the CCW direction than in the CW direction (Fig. 2C and 2D). This result is attributable to the proximity of the short helix and the γ subunit, which maintains the short helix formation (Fig. 5A-5C). In this way, IF_1_ introduces unidirectionality that favors the CW direction.

Note that the unidirectionality is only observed in a torque-applying condition to the γ subunit, where external energy was added to the system. Thus, it does not violate the second law of thermodynamics. Second, we found that the short helix remained intact after the CCW 120° rotation in contrast to the CW 120° rotation. This can be attributed to the reduced change in contact between the short helix and the γ subunit (Fig. 3E and 5B). The short helix may act as a buffer to absorb the γ rotation from external torque. Third, we found that the conformational change of the IF_1_-bound αβ was suppressed in the CCW direction due to the physical blockage of IF_1_ at the αβ interface (Fig. 3A and 3B). It prevents the cooperative catalysis and the subsequent γ rotation. Finally, we identified a steric clash of F22 in IF_1_ with the neighboring β_TP_ subunit. This result agrees with the suggestion from the single-molecule manipulation experiment, where F22 of IF_1_ is crucial for rotation-direction-dependent inhibition and activation. Taken together, these observations indicate that IF_1_ employs various strategies to halt CCW rotation completely, thereby suppressing unfavorable ATP hydrolysis.

### Similarity with other inhibitory systems in ATP synthase

The molecular mechanisms identified in our current study are possibly shared in other inhibitory proteins of ATP synthase. The most similar system to IF_1_ structurally is the ζ subunit for ATP synthase from *Paracoccus dentrificans*. The central long helix in its N-terminus was inserted into the αβ_DP_ interface, while the additional helices in its C-terminus were formed outside of the F_1_ domain (43, 44). The ζ-bound αβ_DP_ interface adopts an intermediate conformation between the loose and tight states, as seen in the *b*MF_1_-IF_1_ complex (Fig. 1A). A prominent difference with IF_1_ is that the ζ does not associate with the γ subunit, as the ζ subunit lacks the amino acid residues corresponding to the residues 1-18 in IF_1_. However, several key residues are conserved, including the F4 residue corresponding to the F22 in IF_1,_ which is crucial for the rotation-direction-dependent regulation, and E12 corresponding to the E30 in IF_1,_ which is crucial for stabilizing the inhibitory complex. Therefore, it is highly probable that the ζ acts as an inhibitory protein in a very similar way to IF_1_. The single-molecule manipulation experiment of the F_1_-ζ complex and the torque-applying simulations, as conducted in our research, would provide more details on the inhibitory mechanism of the ζ subunit. Another example of the inhibitory system is the ε subunit for bacterial ATP synthase, which is sequentially and structurally different from IF_1_. Recent cryo-EM structures of bacterial F_1_ and F_o_F_1_ elucidated that the inhibitory C-terminus of the ε subunit is inserted into the central crevice formed by α_DP_, β_DP,_ and γ, almost in parallel to the γ subunit (45–47). The unique binding mode of ε forces β_DP_ to take an open state with no bound nucleotides, as in the typical αβ_E_ state. These structures suggested a steric clash between ε and the β_TP_ subunit would happen when the γ subunit rotates to the ATP hydrolysis direction because β_TP_ takes the closed conformation with its C-terminal domain located closer to the γ and ε subunits (45, 46). This claim agrees with our findings, although the binding mode differs from IF_1_. Thus, blocking rotations in the ATP hydrolysis direction seems universal among inhibitory proteins. Notably, mycobacterium ATP synthase uses the C-terminal extended region of the α subunit that associates with the globular domain of the γ subunit for its inhibition, which is still much to be explored (48–50). It would be interesting to find similarities with IF_1_, ζ, and ε from the mechanistic point of view.

## Methods

### Structural comparisons of the *b*MF_1_

The 26 structures (78 αβ pairs) of *b*MF_1_ from the following PDB IDs were analyzed by principal component analysis (PCA): 1bmf (51), 1cow (52), 1e1q (53), 1e1r (53), 1e79 (54), 1efr (55), 1h8e (56), 1h8h (56), 1nbm (57), 1ohh (31), 1w0j (58), 1w0k (58), 2ck3 (59), 2jdi (18), 2jiz (60), 2jj1 (60), 2jj2 (60), 2v7q (19), 4asu (61), 4tsf (20), 4tt3 (20), 4yxw (30), 4z1m (30). Considering the resolved residues in all structures, the following residues in the α and β subunits were subjected to analysis: residues 24-401, 413-483, 494-509 of the α subunit, and residues 10-126, 129-310, 312-387, 396-464 of the β subunit, respectively. After aligning the Cα atoms of the αβ pairs, the average structure of all αβ pairs was calculated. Then, we computed the covariance matrix of the Cα positions and diagonalized the covariance matrix to obtain the eigenvalues and the eigenvectors. The first two PCA modes explained 97% of the total motions (Fig. 1B). The analysis was performed using the Python packages MDTraj (62) and Numpy.

The rotary angle of the abovementioned 26 *b*MF_1_ structures was calculated as described in the previous paper. To align all α_3_β_3_-ring structures, the coordinates of the F_1_ structure were translated so that the centers of the three centers of mass (Cα atoms) of the N-terminal domain (residues 10-82) of the β subunit become the origin. Also, F_1_ was rotated so that the z-axis became perpendicular to a surface formed by the three centers of mass of the N-terminal domain of the β subunits. F_1_ was then rotated around the z-axis so that the center of mass of the N-terminal domain of β_DP_ was placed on the x-axis. These procedures were also performed before the MD simulation of the *b*MF_1_-IF_1_ complex to orient the γ subunit parallel to the z-axis. With the aligned α_3_β_3_-ring structures, the γ subunit of each F_1_ was best fitted to that of the 1bmf structure using residues 1-30 and 221-270. Based on the rotation matrix from the best fitting, the rotary angle of the γ subunit was defined by the rotation angle of the x-axis projected on the xy-plane.

### Molecular dynamics (MD) simulation

The MD simulation of the IF_1_-bound bovine mitochondrial F_1_ (*b*MF_1_) was performed based on the previous work with slight modifications. The initial structure was taken from the IF_1_-bound crystal structure of *b*MF_1_ (PDB: 2v7q) (19). Residues 23-510 were used for the α subunits, 9-478 were used for the β subunits, 1-272 were used for the γ subunit, all the residues were used for the δ and the ε subunit, and 1-60 were used for IF_1_. MODELLER (63) was used to model the structure of missing residues. Water molecules in the crystal structure were retained unless they overlapped with replaced ligands. The bound nucleotides of the β_TP_ and the β_DP_ subunits were changed to ATP and ADP + P_i_, respectively, representing the post-hydrolysis state. The bound nucleotides in the three α subunits and Mg^2+^ ions were retained as in the crystal structure.

The modeled *b*MF_1_-IF_1_ complex was solvated with TIP3P water (64) in a rectangular box such that the minimum distance to the edge of the box was 10 Å. Then, 150 mM KCl was added to neutralize the system. The total number of atoms is ∼320,000. The Amber ff14SB force field (65) was used for the protein, with the previously developed parameters for ATP and ADP (66), and for P_i_ (28). The system was energy minimized and equilibrated under isothermal–isobaric (NPT) conditions with Ewald electrostatics and restraints on all heavy atoms in the protein for 500 ps and, subsequently, with restraints on only Cα atoms for one ns. After the equilibration, production runs were performed with restraints on the Cα atoms of the 10 N-terminal residues of each of the three β subunits, mimicking the role of His-tag anchoring in the single-molecule experiments. If necessary, the torque was applied to stall the γ subunit at the initial position before starting the forcible rotation. NAMD 2.12 or 2.14 was used for the MD simulation with periodic boundary conditions (67). Langevin dynamics with 1 ps^-1^ damping coefficient was used for temperature control at 310 K, and the Nose–Hoover Langevin piston was used for pressure control at 1 atm (68). The torque-applying simulations in both CCW and CW directions were performed at the rate of 1°/ns of γ rotation as in the previous studies (27, 28).

### Simulation after 120° state in the CW direction

After the first torque-applying simulation to the 120° state in the CW direction, the additional equilibrium run was performed with the γ subunit coordinates restrained for 100 ns. We then extracted the final snapshot of the simulation and performed the system setup for the next 120° simulation. To reflect the chemical state of the CW 120° state, the bound nucleotides of the β subunits were replaced as follows. The αβ_1_ pair, where ADP and P_i_ were bound at the 0° structure, has a newly bound ATP. The αβ_2_ pair, where an ATP molecule was bound in the 0° structure, has no bound nucleotide in the 120° state. In the αβ_3_ pair, where no nucleotide was bound at 0° structure, ADP and P_i_ were added to the binding site. The subsequent minimization and equilibrations were performed as described above for the first simulation. To induce the appropriate conformational change at 120°, the targeted MD simulation (69) of each αβ pair was performed by NAMD, reducing the RMSD between the current coordinates and the target structure. The target structure for reference was taken from the ground-state structure of *b*MF_1_ (PDB: 2jdi): the αβ_2_ was forced to take the Empty structure, and the αβ_3_ was forced to take the DP structure. This procedure was not performed on the αβ_1_, as an appropriate conformational change of the αβ_1_ was already observed during the forced γ rotation. The equilibrium run was again performed with the γ subunit coordinates restrained for 100 ns.

The second torque-applying simulation was performed from the CW 120° to the CW 240° state. However, crucial changes in IF_1_ were not observed. After the additional equilibrium run at this angle with the γ subunit restrained for 100 ns, we replaced the chemical state with the one of the 240° state. At this angle, the nucleotide state of the β subunits is as follows. The αβ_1_ pair has no bound nucleotide, the αβ_2_ pair has ADP and P_i_, and the αβ_3_ pair has ATP (Fig. 4A). Then, the targeted MD was performed to induce the conformational change of each αβ pair: the αβ_1_ was forced to take the Empty structure, the αβ_2_ was forced to take the DP structure, and the αβ_3_ was forced to take the TP structure, respectively. The abovementioned procedure in the first and the second targeted MD is summarized in the Table. S1.

### Torque-applying simulation

In this study, the rotational angles of the γ-subunit, represented by its Cα atoms, were computed using a displacement vector projected onto a plane perpendicular to the rotation axis. The displacement vector 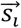 for each residue *i* was defined as

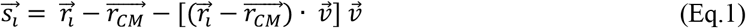

where 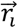 is the positional vector of residue 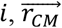 is the center of mass of the Cα atoms, and the 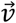 represents the rotation axis (the z-axis in the current study). This projection ensures that the displacement vector lies within the plane perpendicular to the rotation axis. The rotational angle *φ*_*i*_ for each residue *i* was then calculated as the angle between the initial displacement vector 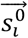 and the current displacement vector 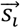,using the following expression:

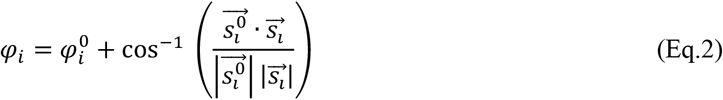

where 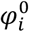 represents the initial rotational angle. To eliminate the overall translation of the γ-subunit, the center of mass 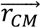 was subtracted from the positional vectors.

In the simulations, distance-weighted averages of the cosine and sine of the rotational angles were controlled. The weight is the length of the displacement vectors 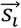.The distance-weighted average cosine and sine of the rotational angle were given by:

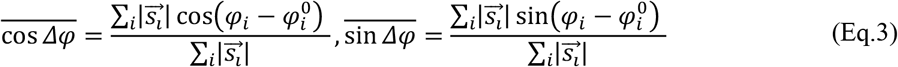

with,

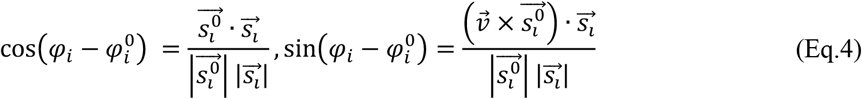

The torque potential *V*(*t*) was defined as:

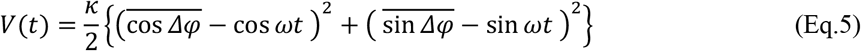

where, κ is the force constant (10^4^, 10^5^ and 10^6^ kcal · mol^-1·^ rad^-2^), *ω* is the angular velocity of the rotation (1 °/ns). For the analysis, the rotation angle of the γ subunit (Fig. 2C and 2D) was calculated as:

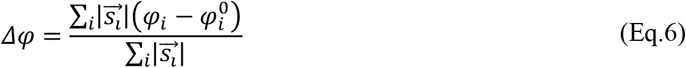

### Work and torque estimation

The non-equilibrium work at time *t* by the torque potential is defined as the previous paper (70):

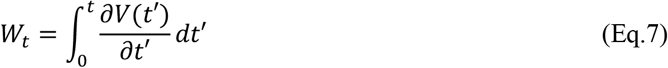

We obtain

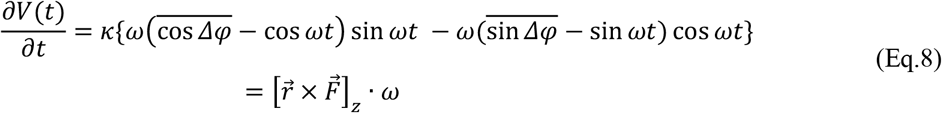

Here, we define

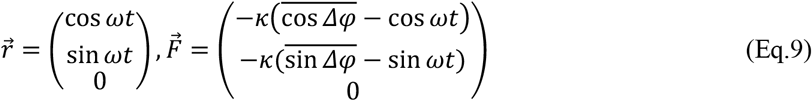

Then,

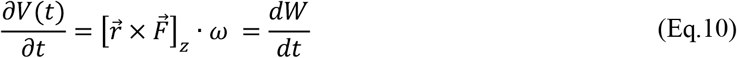

where 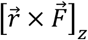 represents the applied torque in the simulations.

## Acknowledgments

We thank Dr. Hiroyuki Noji and Dr. Hiroshi Ueno (University of Tokyo) for the fruitful discussion and all members of the Okazaki group for their valuable comments. This work was partly supported by Grants-in-Aid for JSPS Fellows (JP22KJ3188 to R.K.) and Scientific Research (JP22H02595 and JP23K23858 to K.O.). The computation was partially performed using the Research Center for Computational Science, Okazaki, Japan (Project: 22-IMS-C189 and 23-IMS-C201).

## Author Contributions

R.K. and K.O. designed the research, conducted molecular dynamics simulations and analyses, and wrote the paper.

