## Supplementary material for "Rotation-direction-dependent regulation of ATPase inhibitory factor 1 for mitochondrial ATP synthase from atomistic simulation": SI file

This PDF file includes:

- Figs. S1 to S9
- Table. S1

[1]*Research Center for Computational Science, Institute for Molecular Science, National Institutes of Natural Sciences, Okazaki, Aichi 444-8585, Japan*
[2] *Graduate Institute for Advanced Studies, SOKENDAI, Okazaki, Aichi 444-8585, Japan*

*Corresponding author: Kei-ichi Okazaki

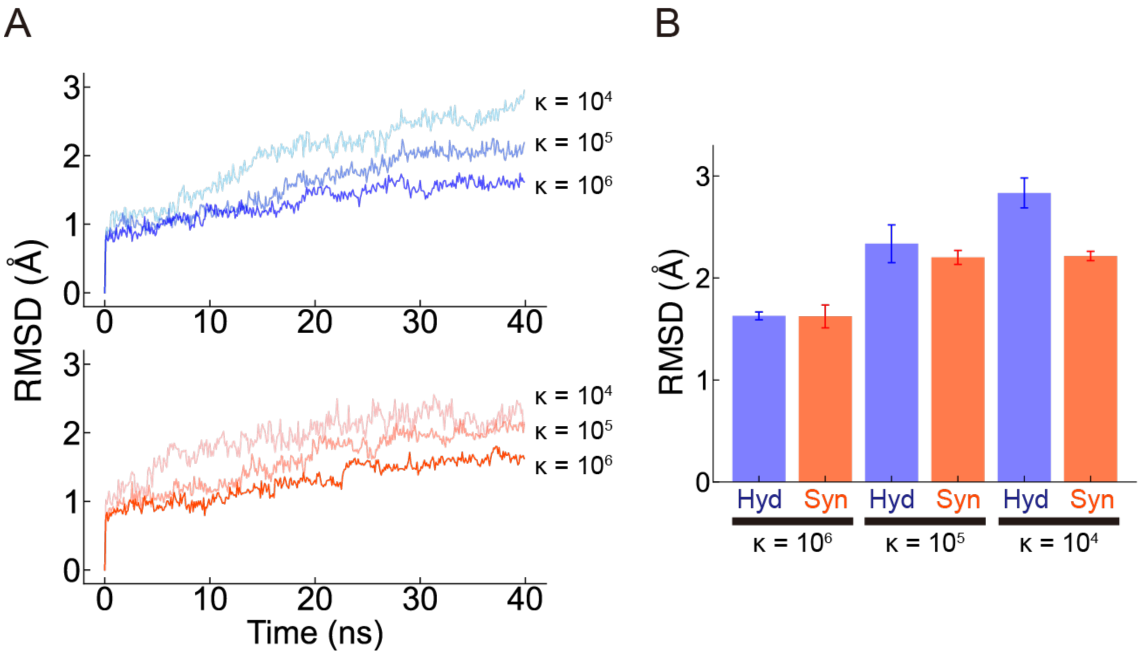


**Fig. S1. Twisting of the γ subunit during the torque-applying simulation.**
(A) Root-mean-square deviation (RMSD) plot during 40 ns simulation of CCW direction (upper, blue) and CW direction (lower, orange). (B) The average RMSD value at 40 ns. Bars represent SD from three independent simulations.

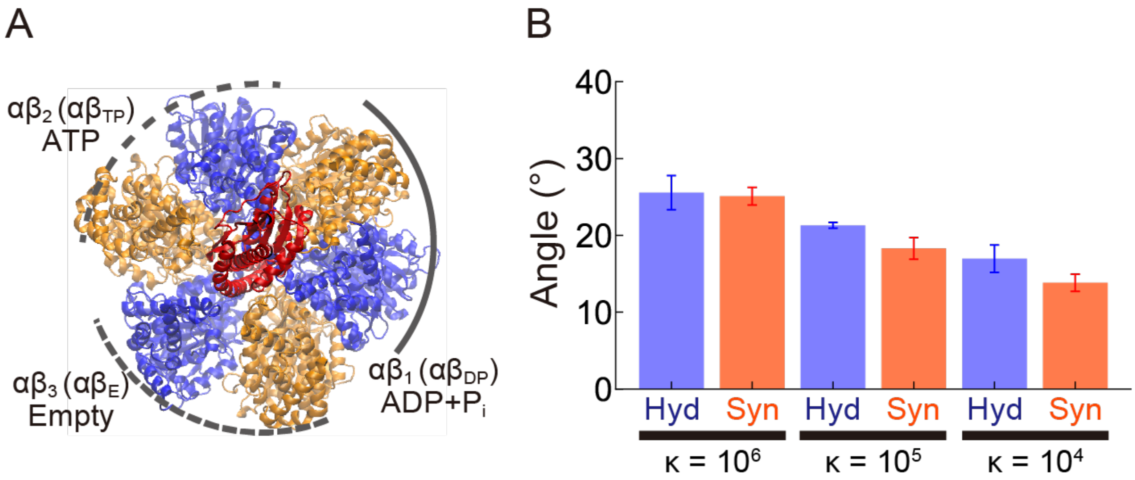


**Fig. S2. Simulations of $IF_1$-free *b*$MF_1$.**
(A) The top view of the $F_1$ structure with the initial bound nucleotides. The δ and ε subunits are omitted in this figure. (B) The final core rotation from the 40 ns simulation. Values represent the mean, and error bars represent SD, estimated from 3 independent simulations.

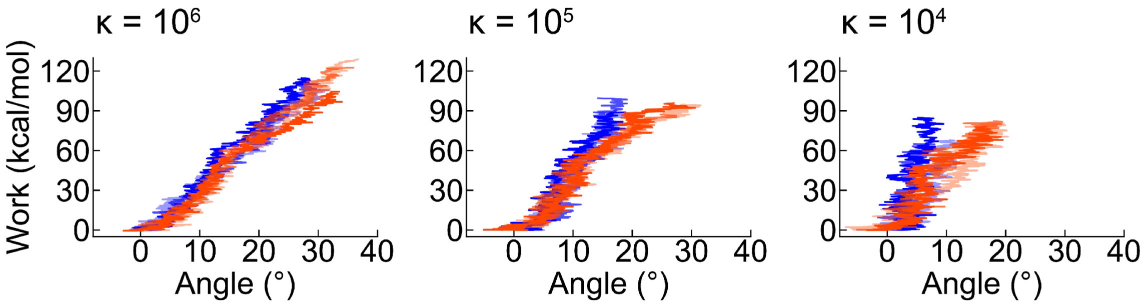


**Fig. S3. Non-equilibrium work along the core angle with three different κ values.**
The blue and red lines represent CCW and CW rotations, respectively. Three simulation trajectories are shown in gradation. The plots show that the increase of non-equilibrium work is steeper for CCW rotation (the hydrolysis direction) than CW rotation (the synthesis direction) along the core angle.

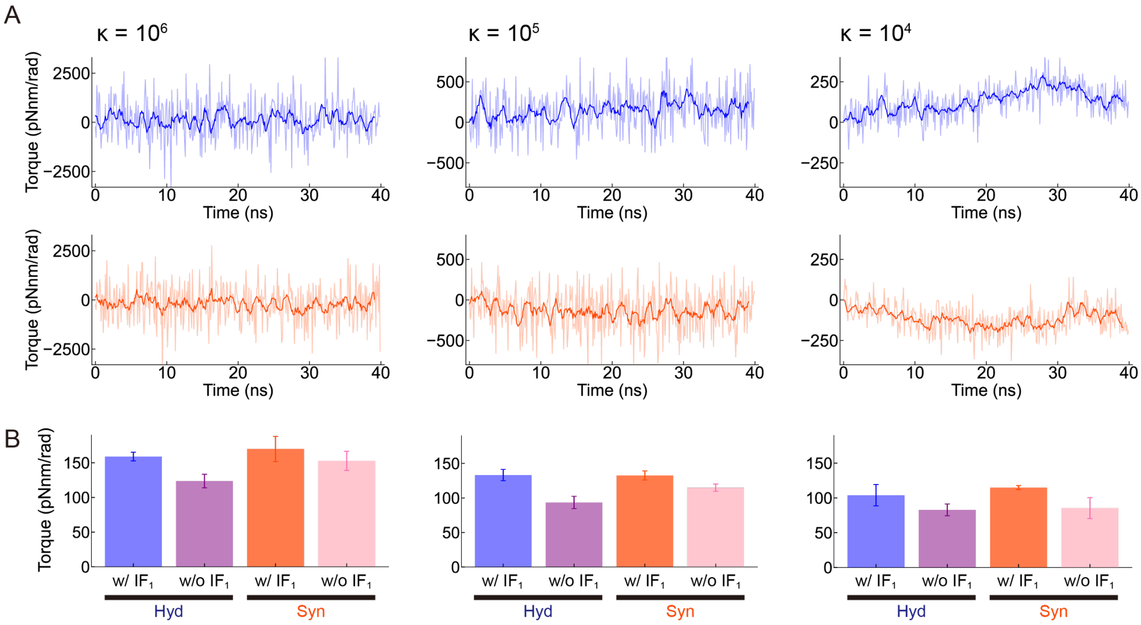


**Fig. S4. Torque estimation during simulations.**

(A) The torque plots of the CCW (*Upper*) and the CW (*Lower*) simulations of the $IF_1$-bound $F_1$. Positive and negative are defined as CCW and CW directions, respectively. The raw data and the running average are represented by light and dark colors, respectively. (B) The average torque values from three independent simulations are plotted. The values in the CW direction were converted to positive ones. Values and error bars represent the mean and the SD, respectively.

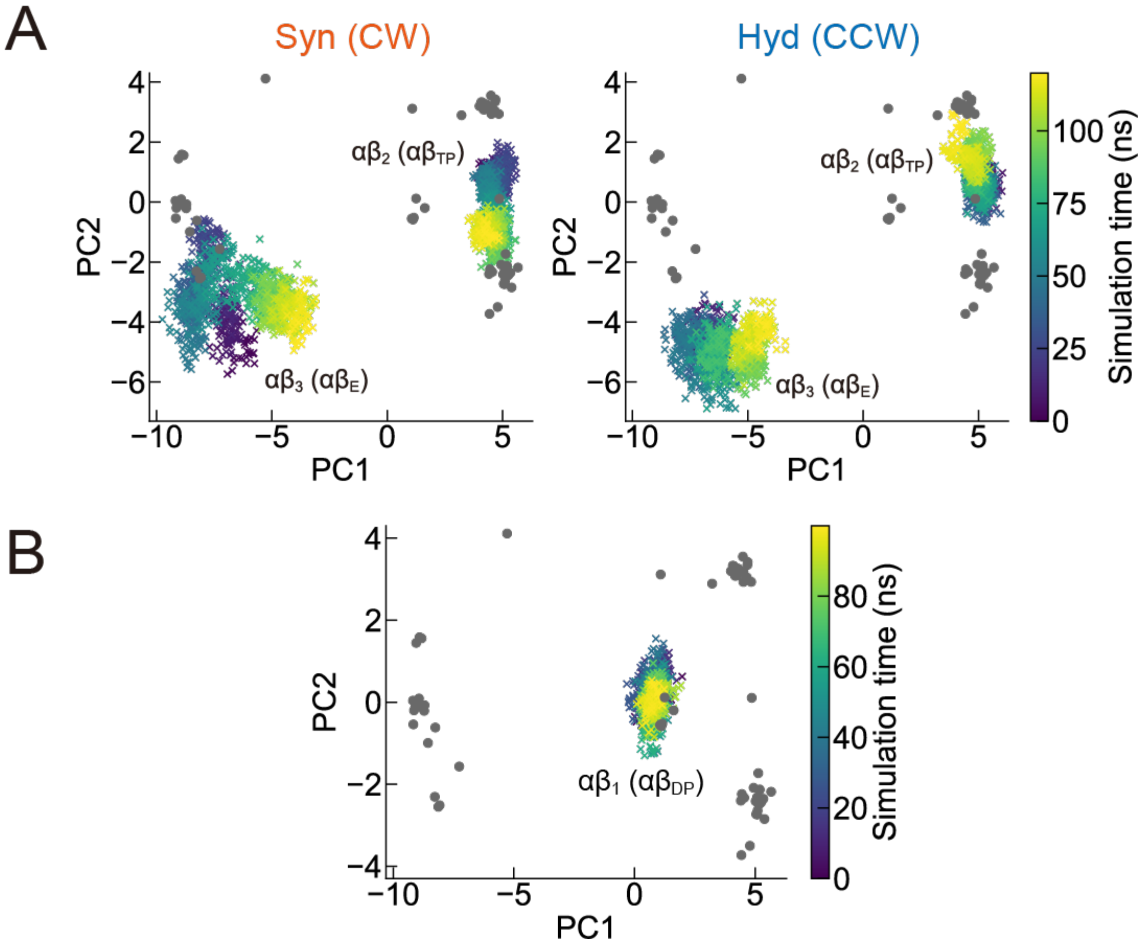


**Fig. S5. Conformational changes of each αβ pair projected onto the PC1-PC2 plane, determined by PCA of the X-ray crystal structures shown in gray dots (Fig. 1B).**

(A) Conformational changes of the $\alpha\beta_2$ ($\alpha\beta_{TP}$) and $\alpha\beta_3$ ($\alpha\beta_E$) pair upon the 120° rotation with the CW (left) and CCW (right), whereas that of the $IF_1$-bound $\alpha\beta_1$($\alpha\beta_{DP}$) is presented in Fig. 3A and 3B. (B) Conformational change of the $IF_1$-bound $\alpha\beta_1$ ($\alpha\beta_{DP}$) during 100 ns of the equilibrium run before the torque-applying simulation.

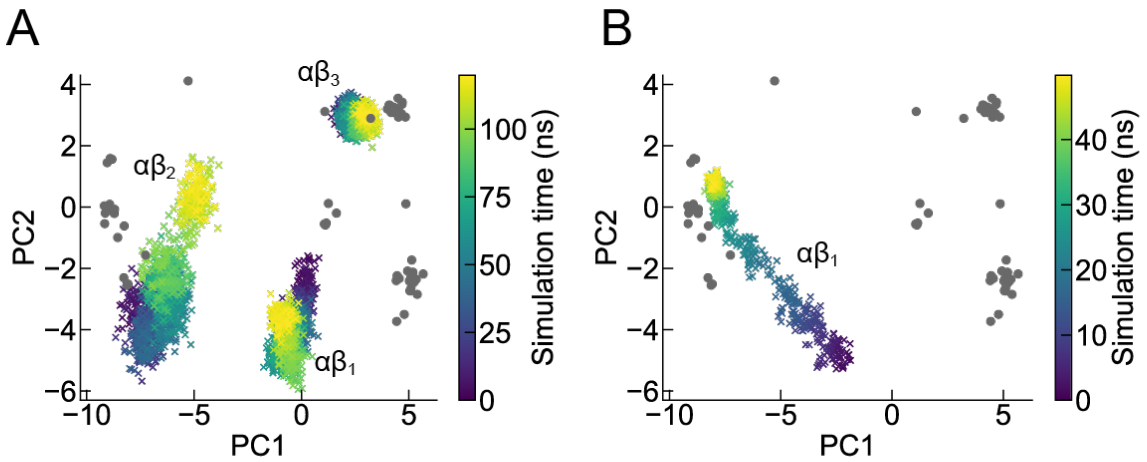


**Fig. S6. Conformational change of the αβ pairs after CW 120° rotation.**
(A) The CW γ rotation from 120° to 240°. (B) The targeted MD at 240°. The gray dots correspond to the X-ray crystal structures (Fig. 1B).

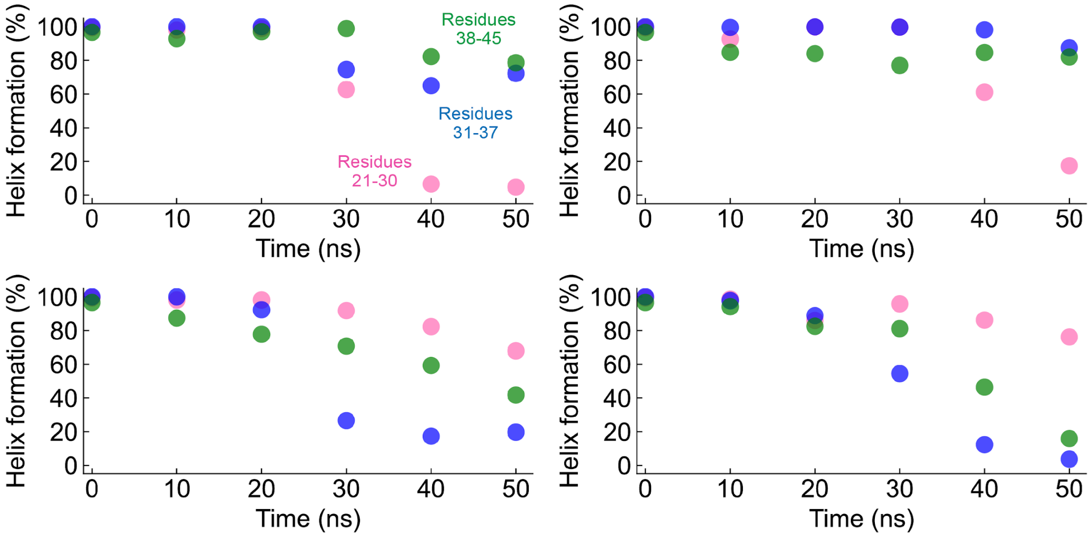


**Fig. S7. Time-dependent long helix deformation in four independent runs during the targeted MD at CW 240°.**

The long helix was divided into three parts: residues 21-30 (pink), 31-37 (blue), and 38-45 (green). The helix formation at time 0 was estimated from the simulation result before the torque-applying simulation. The left upper one is the same as Fig. 4C. The upper two results show that the deformation of the long helix occurred at the entrance of the long helix (pink), while the lower two results show that the deformation occurred at the second half of the long helix (blue).

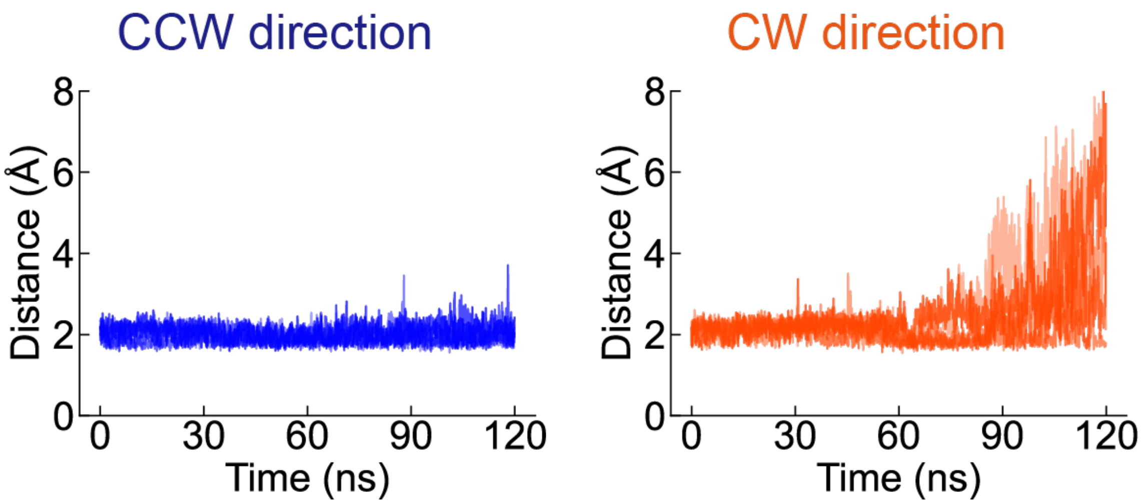


**Fig. S8. Minimum distance between the short helix and the γ subunit during 120° simulations.** Five simulation results were included in different shades of color.

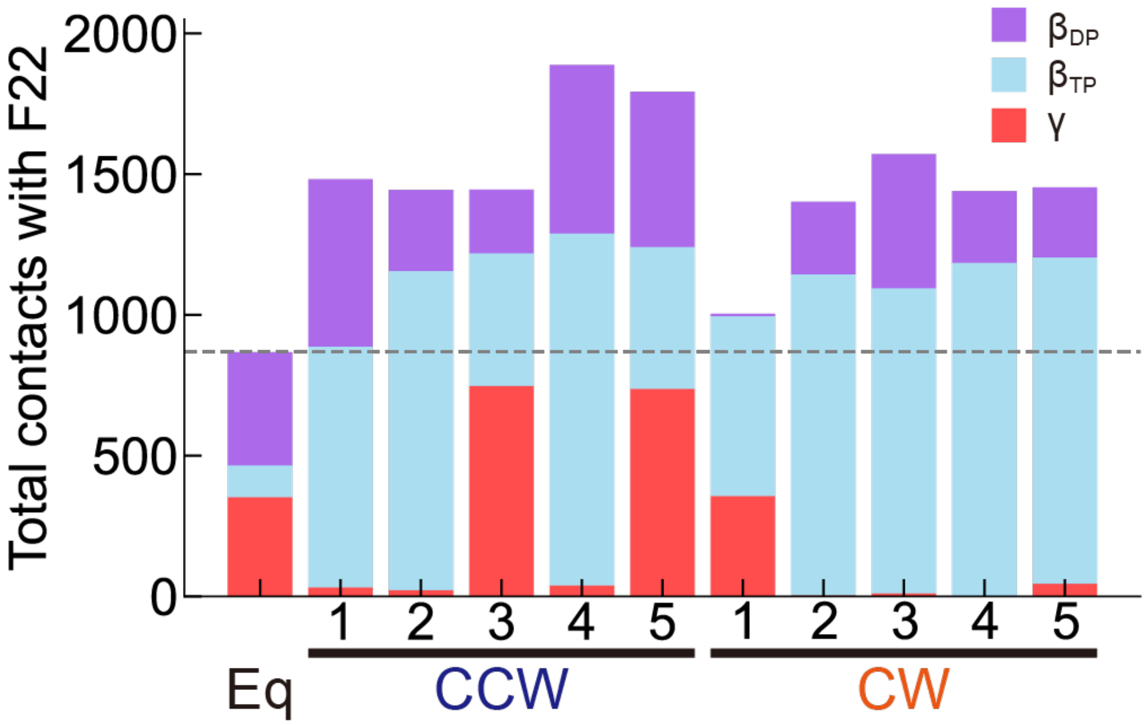


**Fig. S9. Contact analysis of F22 from $IF_1$ during the last 30 ns (300 frames) of each simulation.** The result of ‘Eq’ represents the simulation result before the torque-applying run (= torque-free run). The dotted line represents the total number of contacts in the ‘Eq’ run as a reference value.

**Table. S1. The conformational states and the bound nucleotides in the initial structure, the 1$^{st}$ and 2$^{nd}$ targeted MD.**

The $\alpha\beta_E$, $\alpha\beta_{TP}$, and $\alpha\beta_{DP}$ represent the typical conformation of the αβ pair: $\alpha\beta_E$ adopts an open conformation of the β subunit, and the $\alpha\beta_{TP}$ and $\alpha\beta_{DP}$ adopt a closed conformation with its interface loose or tight, respectively. The bound nucleotides are described in the parentheses.

| | Initial state | 1$^{st}$ targeted MD | 2$^{nd}$ targeted MD |
|---|---|---|---|
| $\alpha\beta_1$ | $\alpha\beta_{DP}$ (ADP + $P_i$) | $\alpha\beta_{TP}$ (ATP) | $\alpha\beta_E$ (Empty) |
| $\alpha\beta_2$ | $\alpha\beta_{TP}$ (ATP) | $\alpha\beta_E$ (Empty) | $\alpha\beta_{DP}$ (ADP + $P_i$) |
| $\alpha\beta_3$ | $\alpha\beta_E$ (Empty) | $\alpha\beta_{DP}$ (ADP + $P_i$) | $\alpha\beta_{TP}$ (ATP) |
